# Identifying Representative Network Motifs for Inferring Higher-order Structure of Biological Networks

**DOI:** 10.1101/411520

**Authors:** Wang Tao, Yadong Wang, Jiajie Peng, Chen Jin

**Author notes:** Equal contributor (Wang Tao, Jiajie Peng).

## Abstract

Network motifs are recurring significant patterns of inter-connections, which are recognized as fundamental units to study the higher-order organizations of networks. However, the principle of selecting representative network motifs for local motif based clustering remains largely unexplored. We present a scalable algorithm called FSM for network motif discovery. FSM accelerates the motif discovery process by effectively reducing the number of times to do subgraph isomorphism labeling. Multiple heuristic optimizations for subgraph enumeration and subgraph classification are also adopted in FSM to further improve its performance. Experimental results show that FSM is more efficient than the compared models on computational efficiency and memory usage. Furthermore, our experiments indicate that large and frequent network motifs may be more appropriate to be selected as the representative network motifs for discovering higher-order organizational structures in biological networks than small or low-frequency network motifs.

## I. INTRODUCTION

Complex systems exhibit rich and diverse higher-order organizational structures and biological organisms are no exception [1], [2]. Studying network structures, esp. the higher-order network connectivity patterns, is critical for elucidating the fundamental mechanisms that control and mediate complex behaviors in a biological system [1], [3], [4]. The most common higher-order structures are small to meso-scale subgraphs of a network, which are referred to as network motifs [1]. Based on known network motifs, tools (such as SNAP and MAPPR) have been developed to study network organization in a number of networks using local higher-order graph clustering [2], [5], [6]. For example, the studies on the *C. elegans* neuronal network and airport network have revealed the information propagation unit and the hub structure [2].

Network motifs have been found in various natural networks such as the networks of biochemistry, neurobiology, and ecology [1], [3]. By definition, network motifs are recurring subgraphs that appear more frequent in a target network than in random networks [7]. A network motif is supposed to be biological or statistical significant [3]. Network motifs with a similar structure are considered as building blocks to provide a specific function (such as information passing) for complex systems [1]. Studying network motifs is effective towards understanding the system of biological interactions locally. However, it remains an open question which network motifs are the most appropriate for local higher-order graph clustering. More precisely, will using larger network motifs lead to improved clustering and knowledge discovery in networks? Or is a sparse network motif more capable of identifying higher-order patterns in peripheral regions of a network? While network motifs have been recognized as fundamental units to analyze complex networks, the principle of network motif selection remains largely unexplored.

To systematically explore the network motif selection problem, we face two computational challenges. First, given a biological network with thousands of vertices, the possible number of network motifs increases exponentially with the increase of network motif size (i.e. the number of vertices) [8]. For example, the total number of size-5 network motifs in the *C. elegans* neuronal network is 3,509, while the number for size-6 network motifs is increased to 66,700. It is practically impossible to exhaustively test all possible network motifs ranging from small to meso-scale for local higher-order graph clustering [6]. Furthermore, a vast number of motif-based clusters would be produced if many network motifs are tested, for which it is difficult to determine which one is the most appropriate. New strategies to optimize the selection of network motifs for inferring higher-order organizations in networks are required.

The second challenge of local higher-order graph clustering lies on the network motif discovery problem itself. Tools have been developed for identifying network motifs [7], [9]–[15]. However, mining network motifs is still a computationally challenging task, provided that it has to enumerate subgraphs and count their frequencies in a target network and in numerous randomized networks. In particular, to match a subgraph instances, the Nauty algorithm [16] is often employed for graph isomorphism testing, of which O(*k*!) is the complexity in the worst case (*k* is the subgraph size) [17]. It is common to spend days to derive small to meso-scale network motifs from biological networks with thousands of vertices [7], [14].

In summary, a holistic solution is clearly needed to identify or select representative network motifs for studying higher-order organizations of biological networks. This work requires to quickly obtain enough number of network motifs and to analyze the motif-dependent organizational patterns using hypothesis-driven approaches.

In this article, we present a new tool called “Fast Scalable Motif” (**FSM**) algorithm, with which, we make three hypotheses on the indentification of representative network motifs and test them systematically. First, since frequent network motifs have more instances in the target network than low frequent ones, they may derive less random clusters. We hypothesize that a representative network motif must be highly frequent (**H1**). Second, the instances of highly similar network motifs may be densely overlapped with each other, leading to highly conserved organizational patterns. We hypothesize that if a organizational pattern is valid, it can be re-revealed by using multiple similar network motifs (**H2**). Third, we hypothesize that clusters derived using large network motifs are larger than the clusters derived using small network motifs (**H3**).

Our contributions are two-fold. First, for a given biological network, it is usually unclear what are the representative network motifs that are most suitable for studying the local higher-order organizations. Only using the existing network motifs identified from other networks will result in significant loss-of-information problem. Since the current network motif discovery tools are computationally prohibitive, esp. for meso-scale network motifs, we present a new method called FSM for efficient network motif discovery. Experimental results show that FSM performs significantly better than the compared methods in both time and memory usage, esp. on meso-scale network motifs.

Second, network motifs in the same biological network could be highly similar to each other regarding graph topology. As a result, their subgraph instances may be densely overlapped with each other, leading to highly conserved organizational patterns. Are the conserved patterns more important than the others? Or they are just redundant information and should be removed? Will using infrequently occurred network motifs result in unreliable higher-order structures due to the lack of subgraph instances? To our knowledge, the organizational network pattern conservation problem has not been extensively studied. Applying FSM on the *C. elegans* neuronal network with 131 vertices and 764 edges [1], we systematically analyzed the clusters obtained with different types of network motifs. The experimental results indicate that *large and frequent network motifs* may be more appropriate to be selected as the representative network motifs for discovering higher-order organizational structures in biological networks.

## II. BACKGROUND

Current studies on biological networks focus on local network properties, i.e. small-world [18], power-law degree distribution [19], network transitivity [20], network motif [1], and community structure [21]. Among them, a widely-used method for exploring higher-order network structures is to identify network motifs using graph mining techniques [7]. In this section, we introduce the network related concepts, followed by the existing methods for network motif discovery and the existing methods of network clustering using network motifs to infer higher-order structures of complex systems.

### A. Network Related Concepts

A large-scale biological network is represented as *G(V, E*), where *V* represents the set of vertices of *G* and *E* = {(*u,v*)|*u* ∈ *V,v* ∈ *V,u* ≠ *v*} represents the edges of *G*. An edge *e* = (*u,v*) also represents the edge direction in directed networks. The vertex degree *deg*(*v*) in a undirected network is defined as *deg(v)* = |{(*v,u*)}|, where *v* ∈ *V*_*u*_ and *u* ∈ *Neighbor*(*v*). The adjacency matrix *A* of network *G(V,E)* is an *n×n* square matrix, where *n* = |*V*| and element *a*_*ij*_ = 1 (or 0) represents there exists (or do not exist) an edge between vertex *v*_*i*_ and *v*_*j*_.

The *canonical form* of a subgraph *g* is the lexicographically smallest or largest adjacency matrix sequence in all permutations of vertex orders. The adjacency matrix that produces the canonical form is called the canonical adjacency matrix (CAM). Two subgraphs *g*(*V*_*g*_, *E*_*g*_) and *h*(*V*_*h*_, *E*_*h*_) are *isomorphic*, if and only if there exists a mapping correspondence between *V*_*g*_ and *V*_*h*_, *f* : *V*_*g*_ → *V*_*h*_, such that (*u, v*) ∈ *E*_*g*_ and (*f(u),f(v)*) ∈ *E*_*h*_. It’s obvious that isomorphic subgraphs have the same CAM and vise versa.

### B. Network Motif Discovery

A number of network motif discovery algorithms have been proposed in recent years [7], [9]–[13], [22]–[25]. FAMMOD takes advantage of a novel algorithm called ESU [26] for subgraph enumeration and utilizes an efficient canonical labeling algorithm called Nauty [16] for CAM calculation [12]. FAMMOD is time-efficient, but it can only handle small network motifs with size less than 9. Kavosh is a memory and time efficient method for network motif discovery in numerically labeled networks [13]. Given vertex *v*_*i*_, it enumerates all ksize subgraph instances containing *v*_*i*_ by implicitly building a tree with *v*_*i*_ as the root and with *k* — 1 levels of descendants based on neighborhood relationship. Kavosh is more efficient than aforementioned models. For example, mining network motifs of size 3 on the *E. coli* network, Kavosh only needs 0.3 seconds while FANMOD and Mfinder need 0.8 seconds and 31 seconds respectively [13]. QuateXelero (QX) is another efficient network motif discovery tool [27]. Similar to FANMOD, it uses ESU for subgraph enumeration and Nauty for canonical labeling. Furthermore, it uses a quaternary tree data structure to store subgraph instances, which can effectively merge instances with the same adjacency matrix and reduce the number of calls of isomorphism labeling (i.e. the most time-consuming step in the whole process). Experiments show that QX achieved better performance than Kavosh. For example, to identify all the size-9 network motifs on the *E. coli* network, QX is 21 times faster than Kavosh [27].

In summary, most existing methods can only detect network motifs with small size (usually less than 10). Furthermore, these methods do not consider network motif characters, such as density and high frequency. In this study, we develop FSM, which is capable of poking our nose into meso or even large-size sparse and highly frequent network motifs, filling the gap between the urgent needs in biomedical research and the currently limited power on network motif discovery approache.

### C. Exploring Network Structures with Network Motif

To understand the higher-order organizational structures of networks at the level of network motifs, Benson *et al*. proposed a spectral clustering method that uses network motifs to cluster the vertices of a network into modules [2]. This method has been integrated into a network analysis package called SNAP [28] to gain new insights of the organization of complex systems. Given a known network motif, SNAP identifies a network motif-dependent organizational pattern, aka a higher-order cluster. SNAP has been successfully applied to reveal the information propagation units in neuronal networks, and hub structure in transportation networks [2].

Network motif-based clustering method is rooted from edge-based clustering. Given network *G*, the aim of edge-based clustering is to find a cluster *S*, which is formed by cutting minimum amount of edges in *G* and including as many as vertices in *S*. The conductance of *S* is defined as:

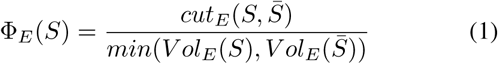

where 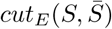indicates the number of edges with one vertex in *S* and the other in 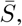 and *Vol*_*E*_(*S*) represents the number of end points in *S*. The lower the conductance is, the better *S* is. Similarly, the network motif based conductance of *S* is defined as:

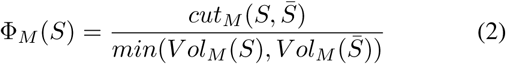

where 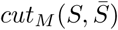 is the number of instances of network motif *M* with at least one vertex in S and one vertex in 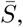 and *Vol*_*M*_(*S*) represents the number of vertices in instances of *M* that reside in *S*. Given network *G(V, E)* and network motif *M*, SNAP identifies *S* with the smallest network motif based conductance using spectral clustering [2], [28].

## III. METHODS

Identifying representative network motifs in a biological network sets the foundation to explore the higher-order organizations of the network. In this section, we introduce a scalable algorithm called FSM for network motif discovery. FSM accelerates network motif discovery by effectively reducing the number of times to do subgraph isomorphism labeling. Multiple heuristic optimizations for subgraph enumeration and subgraph classification are also adopted in FSM to further improve its performance. An illustrative example in Fig. 2 is used to describe all the steps in FSM.

### A. Network Motif Discovery Workflow

FSM consists of three steps, i.e. subgraph enumeration, network motif classification, and significance test. In the workflow, multiple novel strategies of cutting down the computational burden of isomorphism testing are implemented, which make it possible to prioritize network motifs by characters such as frequency and sparseness.

#### Step 1. Subgraph Enumeration

Given a network *G* and a predefined network motif size *k*, we enumerate all the size-k subgraph instances in *G*. There are two enumeration strategies. The first one is the exact enumeration of all possible instances, leading to precise result but with high computational cost. The second strategy is to adopt sampling and statistical approximation, which is faster but less accurate. In this work, we choose the exact subgraph instance enumeration.

#### Step 2. Subgraph Classification

We classify the enumerated subgraphs into isomorphic groups, each corresponding to a unique network motif, represented by CAM. This is the most time-consuming step as it is known to be NP-complete. Nauty [16] is one of the best applications to efficiently generate CAM of a subgraph. FSM adopts Nauty but significantly reduces the number of times of calling Nauty for subgraph isomorphism test compared to the existing methods.

#### Step 3. Significance Test

A list of randomized networks are generated using the original network by keeping the degree distribution. Statistics are performed based on the census to determine whether a subgraph appears significantly more frequent in the original network than in randomized networks.

### B. Step 1. Subgraph Enumeration

FSM adopts an efficient subgraph enumeration method called ESU, first implemented in FANMOD [12]. Mathematically, given a network *G(V, E)* with *N* vertices (see an example in Fig. 2A) and network motif size *k*, the enumeration procedure ESU generates all the size-k induced subgraphs in *N* iterations (see algorithm 1). First, we sort the IDs of all the vertices in *G* from smallest to largest and save the sorted list in queue *Q*_*v*_ = {*v*_1_, *v*_2_,…, *v*_*n*_}. Next, we pop the first vertex from *Q_v_* and choose it as the first vertex *v*_*s*_ of a subgraph-to-enumerate (e.g. *v*_3_ in Fig. 2B). Third, based on the first vertex *v*_*s*_, a breadth-first search on *G*, which also avoids any vertices smaller than *v*_*s*_, is conducted to select the other *k* — 1 vertices of the subgraph (e.g. the leaf nodes in Fig. 2B).

The subgraph enumeration process can be illustrated as constructing a ESU-tree of depth *k*, partially revealed in Fig. 2B. The leaf nodes in an ESU-tree represent all size-k subgraphs starting from the first vertex *v*_*s*_. The intermediate nodes of the tree indicate the recursive enumeration process represented by two vertex subsets *SUB* and *EXT*, where *SUB* is the vertices of the partially finished subgraph (|*SUB*| ≤ *k*) and *EXT* contains the neighbors of any vertices in *SUB*. The partially finished subgraph can be dynamically extended by adding a vertex in *EXT* (see procedure *ExtSubgraph* in algorithm 1). This procedure effectively avoids duplicated enumerations, because each node in the same level of the ESU-tree enumerates exclusive subgraphs. For example, in the second level of the ESU-tree in Fig. 2B, all the subgraphs derived from node {(*v*_3_,*v*_4_), (*v*_5_,*v*_6_, *v*_7_)} contain both *v*_3_ and *v*_4_, whereas subgraphs derived from node {(*v*_3_,*v*_5_), (*v*_6_,*v*_7_)} must contain *v*_3_ and *v*_5_ but not *v*_4_, because *v*_4_ < *v*_5._

Since the ESU-tree based subgraph enumeration process derives densely overlapped subgraphs, a large portion of the enumerated subgraphs may have the identical adjacency matrices. For example, subgraph with vertices {*v*_1_,*v*_3_,*v*_4_,*v*_6_} and subgraph with vertices {*v*_1_,*v*_3_,*v*_4_,*v*_7_} are isomorphic. These subgraphs can be directly merged. For subgraphs with difference adjacency matrices, a traditional approach is to calculate and compare their canonical forms, which is NP-hard in the worst case. We alternatively take advantage of the recursive process to maximally avoid redundant computation. For example, subgraphs induced from {*v*_1_,*v*_3_,*v*_4_,*v*_6_} and {*v*_1_,*v*_3_,*v*_4_,*v*_7_} share vertices {*v*_1_,*v*_3_,*v*_4_}. We only need to compare the edges of *v*_7_ in the graph isomorphic checking. For convenience, we present a series of new concepts in FSM (see examples in Fig. 1B and C).

**Fig.1.**
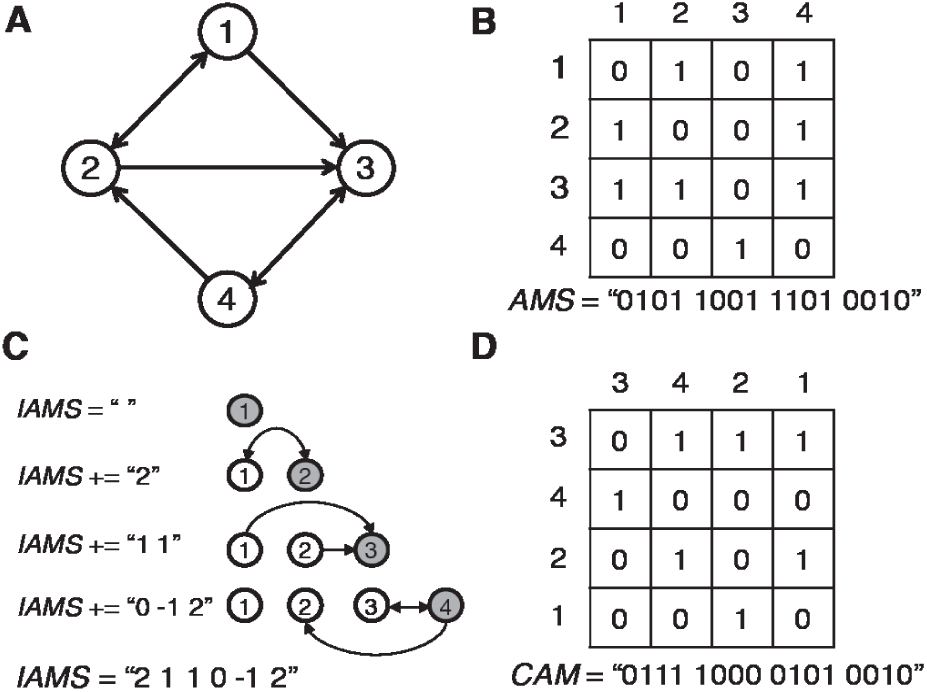
The adjacency matrix string (AMS), iterative adjacency matrix string (IAMS) and canonical adjacency matrix (CAM) of a size-4 subgraph. (A) A directed subgraph. (B) The adjacency matrix of the instance and its corresponding AMS. (C) The subgraph enumeration and IAMS generation process starting with an empty subgraph ***g***. Gray circle represents a new vertex *v_n_* added to *g* in each iteration. For each existing vertex ***vi*** of ***g***, the corresponding value in IAMS has the following meanings:0 for no edge between *v*_*n*_ and *v*_*i*_, 2 for both edges (*v*_*n*_,*v*_*i*_) and (*v*_*i*_,*v*_*n*_), — 1 for edge (*v*_*n*_,*v*_*i*_), and 1 for edge (*v*_*i*_,*v*_*n*_). (D) The CAM of the subgraph in (A).

**Fig.2.**
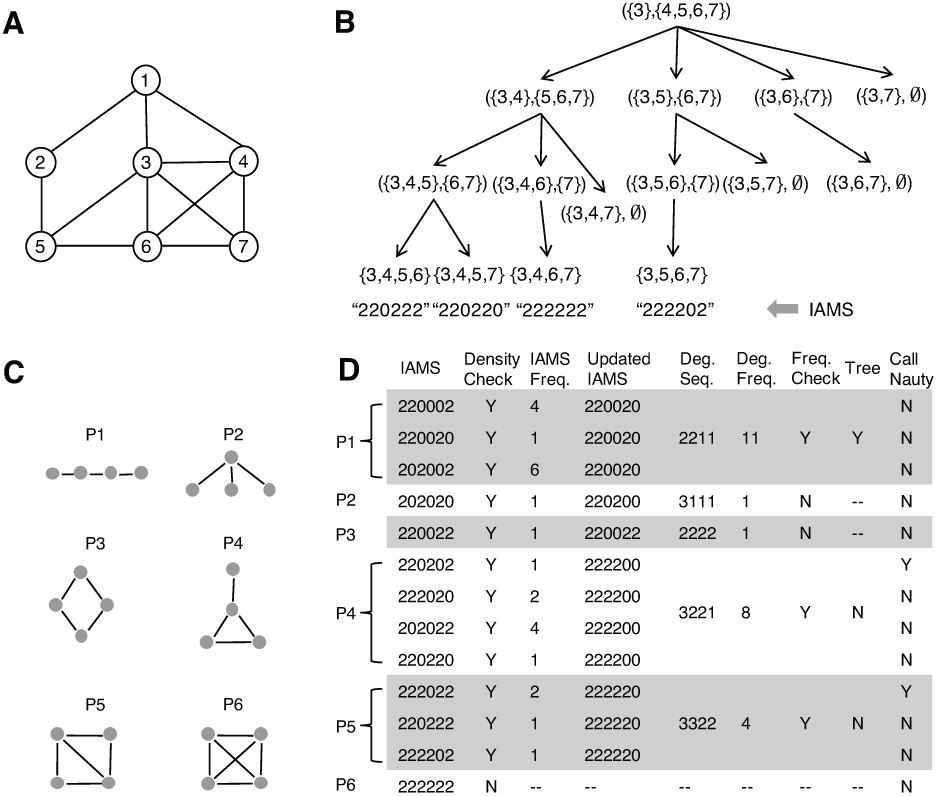
Illustration for subgraph enumeration using ESU and network motif classification. (A) An undirected input network. (B) Steps for enumerating size-4 subgraphs starting from vertex *v*_3_.All the subgraphs are at the leaf level with IAMS labeled beneath. (C) Six non-isomorphic patterns that a size-4 subgraph can form. (D) Steps in network motif classification for reducing the number of times to call Nauty. Isomorphic subgraph groups are separated by pattern labels.

#### Definition III.1. Adjacency Matrix String (AMS).

Given a subgraph *g* with *n* vertices and its *n* x *n* adjacency matrix *A*, the Adjacency Matrix String *AMS(g)* is the binary sequence by concatenating all the entries of *A* in the order *a*_11_ … *a*_1*n*_*a*_21_… *a*_2*n*_ … *a*_*n*1_ … *a*_*nn*_, where *a*_*ij*_ is an element of *A* at the ith row and the *j*th column.

#### Definition III.2. Iterative Adjacency Matrix String

(IAMS). Given a partially complete subgraph *g* with vertex set *SUB*, let *EXT* be the set of neighbors of any vertex in *SUB*, the Iterative Adjacency Matrix String *IAMS(g)* is an iteratively generated string in effect by connecting all the vertices in *SUB* and the new vertices chosen from *EXT*.

#### Definition III.3. Ordered Iterative Adjacency Matrix String (OIAMS).

Given a subgraph-to-enumerate *g* and its iterative adjacency matrix string *IAMS(g)*, the Ordered Iterative Adjacency Matrix String *OIAMS(g)* is the IAMS of *g* with vertices sorted by degree.

Let *g* be a partially complete subgraph subgraph with existing vertex set *SUB*, *v*_*i*_ be a vertex in *SUB*, and *v*_*n*_ be a new vertex chosen from *EXT*. We extend *g* by adding *v*_*n*_ to *SUB* and append *IAMS(g)* with a new vector indicating the edge direction between *v*_*n*_ and all the existing vertices in *SUB*. In the new vector, 1 means an edge pointing from *v*_*i*_ to *v*_*n*_; −1 means an edge pointing from *v*_*n*_ to *v*_*i*_; 2 means a bi-direction edge between *v*_*i*_ and *v*_*n*_, and 0 means there does not exist an edge between *v*_*i*_ and *v*_*n*_ (see Fig. 1C).

Based on the definition of IAMS, a CAM of a subgraph can be represented using the form of IAMS. In the example in Fig. 1D, all the 26 size-4 subgraphs are grouped into 13 return *IAMSH* IAMS classes. Subsequently, at least 50% of times to call Nauty for isomorphism check is avoided in FSM.

~~~
**Algorithm 1** Subgraph Enumeration
**Input:**
 *G(V, E):* Input network
 *k:* Subgraph size
 *St:* Sparseness threshold
**Output:**
 *IAMSH:* Hash table for iterative adjacency matrix strings
1:**procedure** SUBGRAPHENUMERATE
2:      initialize *IAMSH* and *IAMSA*
3: **for all** *v* ∈ *V* **do**
4:     *V*_*sub*_ ⇐{*V*}
5:     *V*_*ext*_ ⇐ {*u*} *V*:*u*>*v*}
6:     *IAMSA*[0] ⇐ *Null*
7:             EXTSUBGRAPH(*V*_*sub*_,*V*_*ext*_,*v,IAMSA*)
    **return** *IAMSH*
8: **procedure** EXTSUBGRAPH(*V*_*sub*_,*V*_*ext*_,*v,IAMSA*)
9   *size* ⇐| *V*_*sub*_|
10:   **if** *size* == *k* **and** sparseness(*G*_*V subg*_) < *St* **then**
11:  *iams* ⇐ Concatenate *k* elements in *IAMSA*
12:  *IAMSH* [*iams*] += 1
13: **else**
14:    **while** *V*_*ext*_ ≠ ∅ **do**
15:      *w* ⇐ Randomly choose a vertex in *V*_*ext*_
16:      *V*_*ext*_ ⇐ *V*_*ext*_ {*w*}
17:      *IAMSA*[*size*] (IAMS(*w*, *V*_*sub*_)
18:      *V*′ ⇐ {*u*∈ *V* : *u* > *v* excluding *w* and *V*_*sub*_}
19:      *V*′ *V*_*ext* U*V*_′
20:             EXTSUBGRAPH(*V*_*sub*_ U{*w*},*V*′ ,*v,IAMSA*)
         **return** *IAMSH*
~~~

Biological networks, such as the protein-protein interaction networks, are usually sparsely connected. To study the topological structures without involving the hub nodes, it is necessary to derive sparse network motifs especially for large motif sizes. In algorithm 1 line 10, a user defined sparseness threshold St is set to prune dense subgraphs, which in turn also significantly reduce the computational time. For example, in Fig. 2D, the last group is excluded from downstream analysis because of its low subgraph sparseness.

### C. Subgraph Classification

Subgraphs with identical IAMS are isomorphic to each other and thus are grouped together for frequency counting. However, IAMS is dependent on the vertex selection order of *EXT* in the enumeration process, which, according to algorithm 1, is random. To further improve the performance of FSM, we reform IAMS by permuting subgraph vertices according to their degree order (see algorithm 2 line 1-4), and define it as the ordered iterative adjacency matrix string (OIAMS, see Definition III.3). In the example in Fig. 2D, 12 IAMS classes are grouped into 5 OIAMS classes.

Using infrequent occurred network motifs for high-order organization discovery may result in unrepresentative clusters due to the lack of enough instances in the target network. To address this issue, we further group OIAMS classes by their degree sequences (see Algorithm 2 line 5-6 and examples in the fifth column of Fig. 2D). If the number of instances of a degree sequence group is less than a user given frequency threshold *Ft*, the group will be discarded (see Algorithm 2 line 7-9). The rationale is that all the isomorphic subgraphs must be in the same degree sequence group, while the opposite is not always true. Therefore, if the frequency of a degree sequence group is less than Ft, then none of the subgraphs in the group can satisfy Ft (proof omitted). For example, let Ft = 4 in Fig. 2D, all the subgraph instances within the degree sequence group ‘3111’ should be discarded because the group only contain one instance.

~~~
**Algorithm 2** Subgraph Enumeration
**Input:**
  *IAMSH:* Hash table for iterative adjacency matrix strings
  *k:* Subgraph size
  *Ft:* Frequency threshold
**Output:**
 *FreqCAM:* Frequent size-k subgraphs
1: **for all** *iams* ∈ *IAMSH* **do**
2:           Calculate degree of each vertex
3:   *oiams* ←  reform *iams*
4:   *OIAMSH[oiams]* += *IAMSH[iams]*
5:   *dseq* ← DegreeSeq(*oiams*)
6:   *DegSeqH[dseq].add(oiams)*
7:**for all** *dseq* ∈ *DegSeqH* **do**
8:   *count* ← Frequency(*DegSeqH [dseq]*)
9:   **if** *count* ≥ *Ft* **then**
10:        *dseqSum* ← DegreeSum(*dseq*)
11:        **if** *dseqSum* = 2x(*k* — 1) **then**
12:          **for all** *oiams* ∈ *DegSeqH[dseq]* **do**
13:    *cam*   ∈ TreeCanonicalLabel(*oiams*)
14:    *CAMH[cam]* +=*OIAMSH[oiams]*
15:   **else**
16:   **for all** *oiams* ∈ *DegSeqH[dseq]* **do**
17:    *cam* ← Nauty(*oiams*)
18:    *CAMH[cam]* += *OIAMSH[oiams]*
19:**for all** *cam* ∈ *CAMH* **do**
20:     **if** *CAMH[cam]* ≥ *Ft* **then**
21:    *FreqCAM[cam]* ← *CAMH[cam]*
  **return** *FreqCAM*
~~~

Given a set of sparse subgraphs, chances are that a portion of them are trees. Since the time complexity of tree isomorphic testing is polynomial [29], we separate trees from subgraphs further reducing the number of times for graph isomorphism labeling (see Algorithm 2 line 7-18). As illustrated in Fig. 2D, FSM only needs to call Nauty twice, compared to call Nauty 26 times in other methods.

### D. Significance Test

While frequent subgraphs can be used directly for multiple purpose, significance test has to carried out for network motif discovery. Traditionally, the Z-score is used to prioritize the uniqueness of network motifs.

## IV. EXPERIMENTAL RESULTS

In the experiment, we compared FSM with QuateXelero (QX), one of the state-of-the-art network motif identification methods on three biological networks. QX is among the best in both time cost and in memory cost [27]. For fair comparison, both FSM and QX were executed on the same computer resource. Both FSM and QX were tested on three mediate to large-scale biological networks: 1) YeastPPI: protein-protein interaction network of budding Yeast [30], 2) HuRI: the human reference protein interactome [31], and 3) String: an integrated human gene interaction network with combined score greater than 0.95 from multiple sources including text mining from publications, experimental data sources, and gene co-expression [32] (see Table 1).

**TABLE I.**
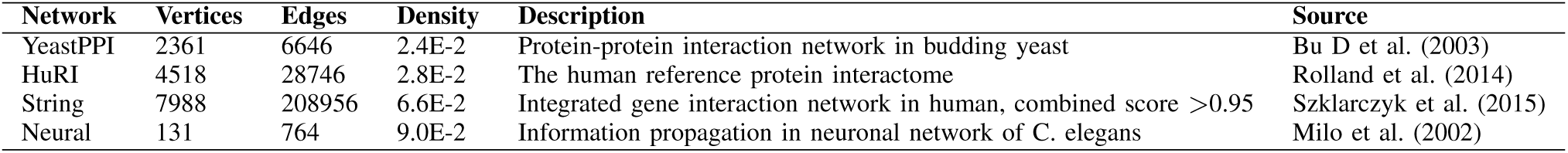
EXPERIMENTAL DATASETS.

### A. Performance of FSM

To discover network motifs with size ranging from 10 to 60, we first enumerated 10 million subgraph instances for each size-*k* subgraph (*k* ∈ {10,20,30,40,50,60}) at each biological networks. Frequency threshold *Ft* was set to 50000 for all network motif sizes, and spareness threshold *St* (the ratio of number of edges and number of nodes) was 2, 4, and 4 in YeastPPI, String and HuRI networks respectively. We compared FSM and QX on the running time, memory, and the number of times of calling Nauty for the subgraph isomorphism test. Note that since both FSM and QX adopt ESU for subgraph enumeration, the subgraph instances mined by both algorithms are identical.

The overall performance is shown in Fig. 3. The first bar plot shows the total number of times of calling Nauty on each of the three tested networks for subgraph canonical labeling. The number of calls made by FSM is significantly less than QX in all the networks (p values < 0.05 by paired t-test). The lower two panels of figure 3 shows that FSM performs better than QX in time and memory esp. on large size network motifs. This is because the extra computational cost in FSM, such as computing IAMS and degree sequences, is significantly less than the cost of calling Nauty, and the difference of the two increase exponentially with subgraph size.

**Fig.3.**
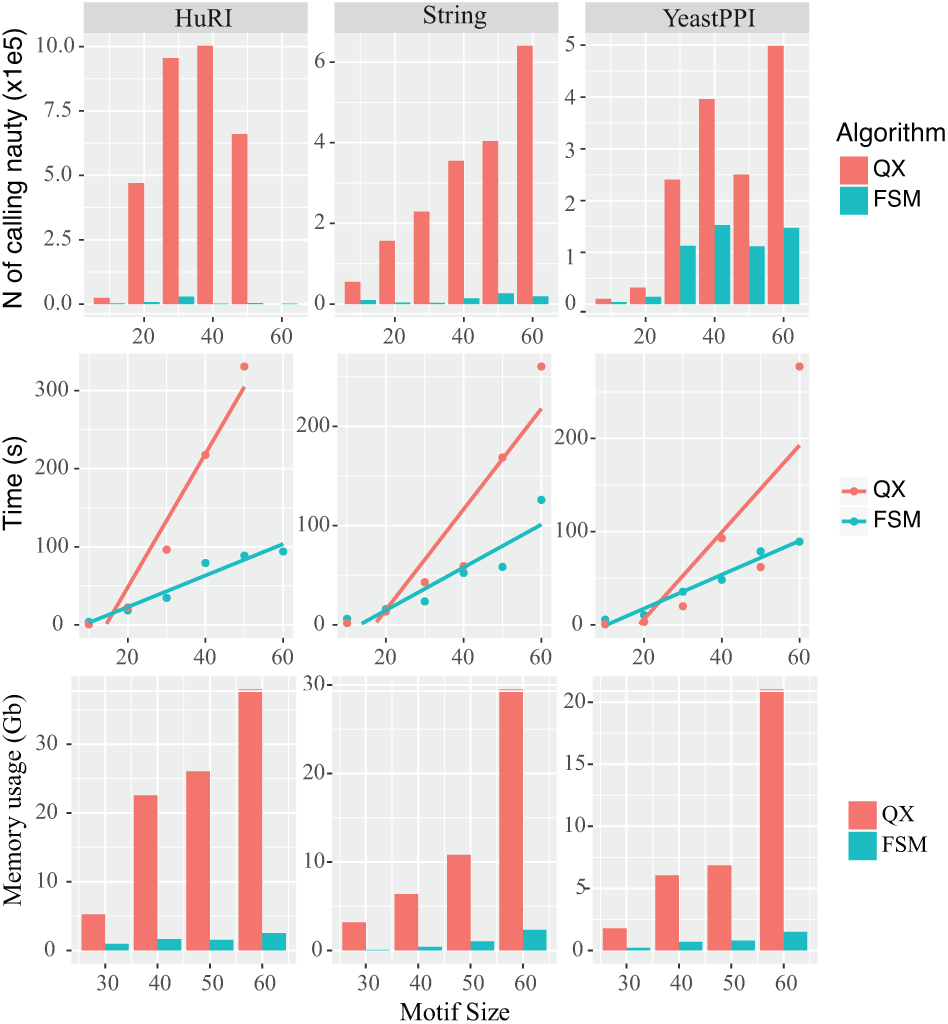
Performance comparison between FSM and QX on network motif discovery on three biological networks.

### B. Higher-order Organization Discovery

Mining network motifs is critical for revealing higher-order organizations of real-world complex networks. With the mined network motifs, SNAP is often employed to gain new insights into the network organization by discovering clusters based on network motifs [2]. However, it remains an open question what are the representative network motifs that are appropriate for local higher-order graph clustering. In a biological network, there could be thousands of network motifs. New strategies are needed to quickly identify local higher-order organizational structures with a few tries. To address this issue and to evaluate our hypotheses, a higher-order cluster detection experiment was performed using all size-5 network motifs on the *C. elegans* neuronal network [1]. Among all the 3,491 subgraphs identified by FSM, we selected 4 groups of subgraphs with different frequency levels for further analysis, i.e. group 1 with 10k-50k instances, group 2 with 1k-5k instances, group 3 with 100-500 instances, and group 4 with only 10-50 instances. We then employed SNAP to discover local higher-order clusters using every subgraph [2]. Note that motif size was limited to 5 because of the small size of the *C. elegans* neuronal network. Our method can be easily extended to discover motif with larger size (see Fig. 3).

Fig. 4 left panel shows the percentages of cluster size (cluster size divided by whole network size) of the four groups of network motifs are significantly different (p value <2e—16, one-way ANOVA). It also reveals that more frequent a network motif is, larger cluster it tends to derive. Finally, the red horizontal line indicates the size of the cluster found using the bifan motif (size-4), much smaller than the clusters found using frequent and large network motifs [2]. Fig. 4 right panel shows the percentage of overlap between the bifan derived cluster and the clusters derived using each network motif in the 4 testing groups, revealing that using frequent network motifs turn to generate clusters that have higher overlap rates with the bifan-derived cluster.

**Fig.4.**
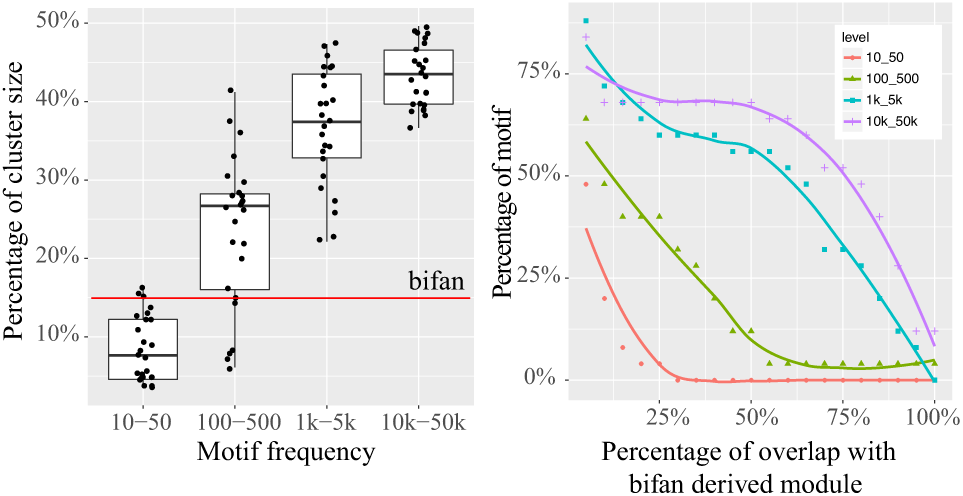
Using frequent network motifs derived larger clusters in *C. elegans* neuronal network. (A) Cluster size distribution of the 4 network motif groups. The red line indicates the size of the cluster derived using bifan. (B) Cluster overlap rates with the cluster derived using bifan.

Next, we tested the conservativeness of the local clusters identified using different network motifs. Conservativeness is defined as the normalized size of the overlapped portion of two clusters *C*_*i*_ and *C*_*j*_ in Equation 3:

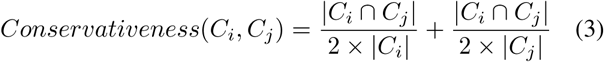

The conservativeness scores among all the clusters derived by the 4 groups of network motifs were visualized as a heat map shown in Fig. 5. The cell color indicates the similarity level between two clusters in corresponding row and column, with red representing higher similarity. The heat map reveals that frequent network motifs (frequency between 10K and 50K) derives highly conserved clusters, while low-frequency network motifs turn to generate non-conservative clusters, which appear to be random.

**Fig.5.**
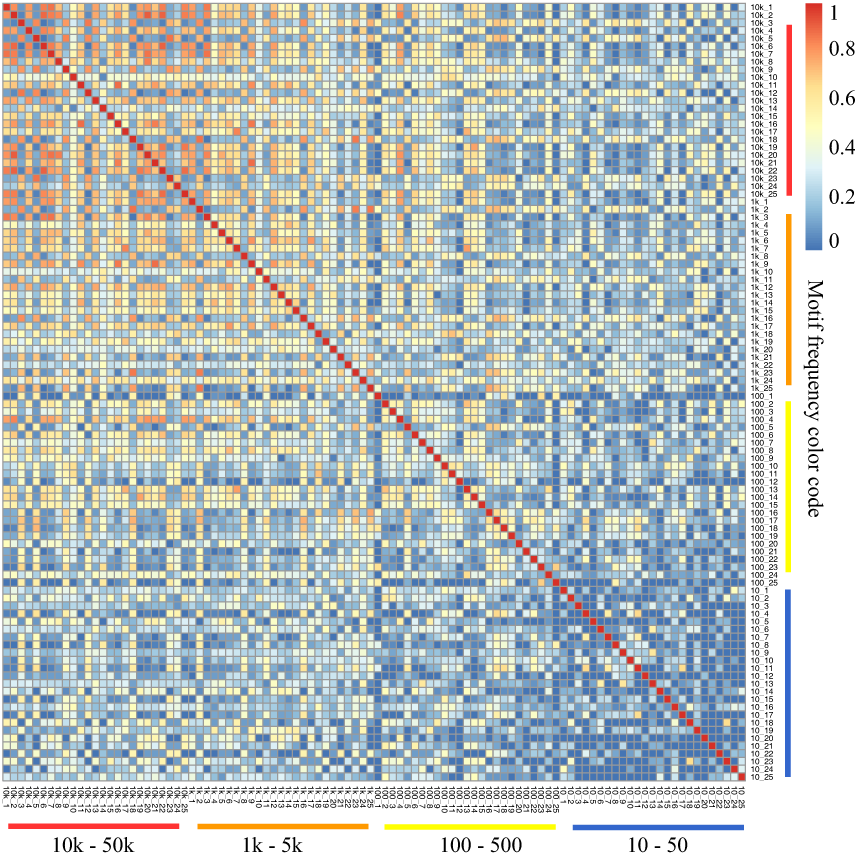
Conservativeness of clusters derived using network motifs with different levels of frequency.

Finally, we test whether the iologically significant bifan-derived cluster [2] is a subset of the highly conserved clusters; and if so, what are the other neurons in the same cluster. Fig. 6A shows a consensus cluster, which is the overlap of the clusters derived using three frequent network motifs shown in Fig. 6B. The red vertices in the consensus cluster are neurons in the bifan-derived cluster, while the blue ones are neurons found using larger network motifs. The consensus cluster can be divided into five sub-modules. Our results contains more neurons in all the five sub-modules, for which, biological significance were found. For example, all the new neurons in the left-most sub-module are ring motor neurons. The bifan-derived cluster includes three out of four RME neurons (i.e. RMEL, RMER, and RMEV). While we found all of them, we also found RMED that was lost by the bifan-derived cluster [33]. The neurons in the top sub-cluster are all sensory neurons of lateral sensilla, and the neurons in the bottom sub-cluster dominated by interneurons [33].

**Fig.6.**
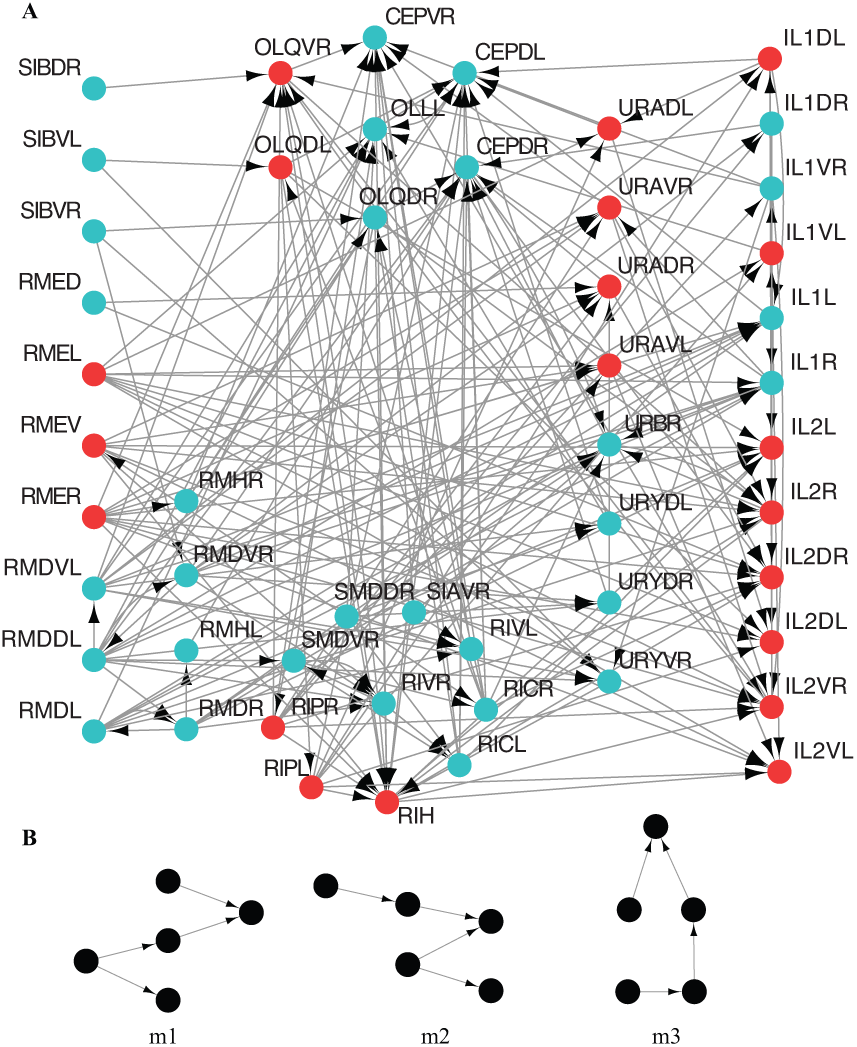
A consensus cluster (Panel A) was identified as the overlap of the clusters derived using three frequent network motifs (Panel B).

## V. DISCUSSION AND CONCLUSION

To elucidate fundamental mechanisms of complex network systems, a computational framework to identify network motifs and then mine clusters in a network by detecting network motif based connectivity has been developed [2]. However, systematic investigation of higher-order organization requires to identify the representative network motifs for any given networks, which has not been well studied in literature. In this article, we propose a novel method FSM to efficiently discover frequent subgraphs and their instances in a given network, and explore the strategies for representative network motif selection. Our experiments on real-world data indicate that *large and frequent network motifs* may be more appropriate to be selected as the representative network motifs for discovering higher-order organizational structures in biological networks than small or low-frequency network motifs.

Our discovery in turn may accelerate network motif discovery process. For example, in the *C. elegans* neuronal network, the frequency distribution and the density distribution of all the size-5 subgraphs are shown in Fig. 7. About half of them have only less than 10 instances, and only a small portion have more than 1,000 instances. Meanwhile, most of the network motifs are sparse. According to our discovery, we can safely remove more than half of the subgraphs without adversely affect the followed network analysis.

**Fig.7.**
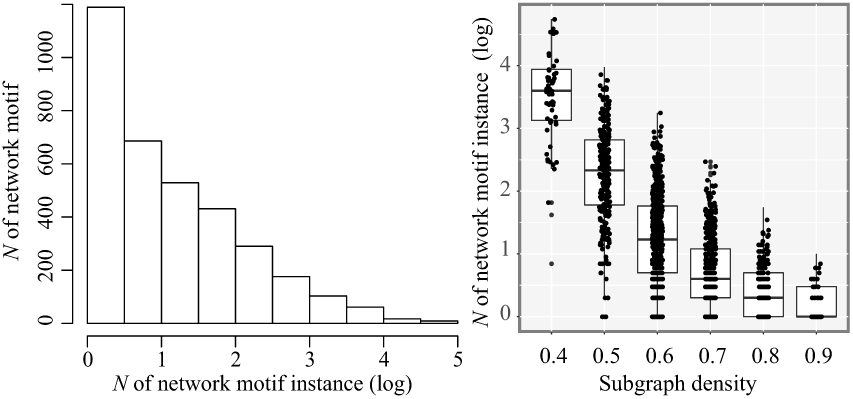
Frequency distribution of all the size-5 subgraphs in *C. elegans* neuronal network.

Using the same frequency threshold for all the network motifs may not be ideal, for the number of subgraph instances varies dramatically when subgraph size increases. An alternative approach is to dynamically adjust the subgraph frequency threshold proportional to the size of the largest bin of OIAMS. Fig. 8 shows the performance on the YeastPPI network after adopting soft thresholding. The coefficient of variation with soft thresholding is 0.55, much lower than 0.75 when a fixed threshold was used, suggesting that soft thresholding may be more reasonable than fixed thresholding on obtaining reasonable number of network motifs for downstream applications. Meanwhile, FSM still performs better than QX on both number of times calling Nauty and on time. In the future, we will incorporate soft-thresholding and systematically study network higher-order structures using large-size network motifs.

**Fig.8.**
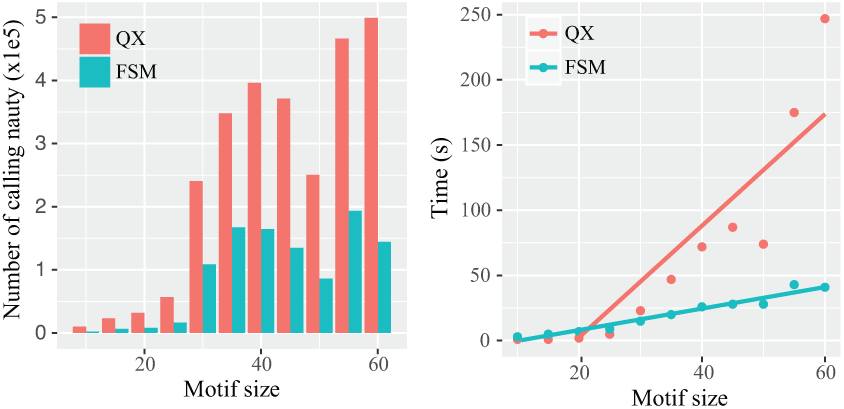
Performance comparison between FSM and QX in YeastPPI network after adopting soft frequency thresholding.

## REFERENCES

[1] R. Milo, S. Shen-Orr, S. Itzkovitz, N. Kashtan, D. Chklovskii, and U. Alon, “Network motifs: simple building blocks of complex networks,” Science, vol. 298, no. 5594, pp. 824–827, 2002.

[2] A. R. Benson, D. F. Gleich, and J. Leskovec, “Higher-order organization of complex networks,” Science, vol. 353, no. 6295, pp. 163–166, 2016.

[3] S. S. Shen-Orr, R. Milo, S. Mangan, and U. Alon, “Network motifs in the transcriptional regulation network of escherichia coli,” Nature genetics, vol. 31, no. 1, p. 64, 2002.

[4] J. Yang and J. Leskovec, “Overlapping communities explain core-periphery organization of networks.” Proceedings of the IEEE, vol. 102, no. 12, pp. 1892–1902, 2014.

[5] Ö. N. Yaveroğlu, N. Malod-Dognin, D. Davis, Z. Levnajic, V. Janjic, R. Karapandza, A. Stojmirovic, and N. Pržulj, “Revealing the hidden language of complex networks,” Scientific reports, vol. 4, p. 4547, 2014.

[6] H. Yin, A. R. Benson, J. Leskovec, and D. F. Gleich, “Local higher-order graph clustering,” in Proceedings of the 23rd ACM SIGKDD International Conference on Knowledge Discovery and Data Mining. ACM, 2017, pp. 555–564.

[7] J. Chen, W. Hsu, M. L. Lee, and S.-K. Ng, “Nemofinder: Dissecting genome-wide protein-protein interactions with meso-scale network motifs,” in Proceedings of the 12th ACM SIGKDD international conference on Knowledge discovery and data mining. ACM, 2006, pp. 106–115.

[8] J. R. Ullmann, “An algorithm for subgraph isomorphism,” Journal of the ACM (JACM), vol. 23, no. 1, pp. 31–42, 1976.

[9] N. Kashtan, S. Itzkovitz, R. Milo, and U. Alon, “Efficient sampling algorithm for estimating subgraph concentrations and detecting network motifs,” Bioinformatics, vol. 20, no. 11, pp. 1746–1758, 2004.

[10] F. Schreiber and H. Schwöbbermeyer, “Mavisto: a tool for the exploration of network motifs,” Bioinformatics, vol. 21, no. 17, pp. 3572–3574, 2005.

[11] P. Ribeiro and F. Silva, “G-tries: an efficient data structure for discovering network motifs,” in Proceedings of the 2010 ACM Symposium on Applied Computing. ACM, 2010, pp. 1559–1566.

[12] S. Wernicke and F. Rasche, “Fanmod: a tool for fast network motif detection,” Bioinformatics, vol. 22, no. 9, pp. 1152–1153, 2006.

[13] Z. R. M. Kashani, H. Ahrabian, E. Elahi, A. Nowzari-Dalini, E. S. Ansari, S. Asadi, S. Mohammadi, F. Schreiber, and A. Masoudi-Nejad, “Kavosh: a new algorithm for finding network motifs,” BMC bioinformatics, vol. 10, no. 1, p. 318, 2009.

[14] W. Lin, X. Xiao, X. Xie, and X.-L. Li, “Network motif discovery: A gpu approach,” IEEE Transactions on Knowledge and Data Engineering, vol. 29, no. 3, pp. 513–528, 2017.

[15] J. Luo, L. Ding, C. Liang, and N. H. Tu, “An efficient network motif discovery approach for co-regulatory networks,” IEEE Access, vol. 6, pp. 14 151–14 158, 2018.

[16] B. D. McKay et al., “Practical graph isomorphism,” 1981.

[17] A. Masoudi-Nejad, M. Ansariola, Z. R. M. Kashani, A. Salehzadeh-Yazdi, and S. Khakabimamaghani, “Cytokavosh: a cytoscape plug-in for finding network motifs in large biological networks,” PLoS One, vol. 7, no. 8, p. e43287, 2012.

[18] L. A. N. Amaral, A. Scala, M. Barthelemy, and H. E. Stanley, “Classes of small-world networks,” Proceedings of the national academy of sciences, vol. 97, no. 21, pp. 11 149–11 152, 2000.

[19] E. Ravasz, A. L. Somera, D. A. Mongru, Z. N. Oltvai, and A.-L. Barabási, “Hierarchical organization of modularity in metabolic networks,” science, vol. 297, no. 5586, pp. 1551–1555, 2002.

[20] M. E. Newman, S. H. Strogatz, and D. J. Watts, “Random graphs with arbitrary degree distributions and their applications,” Physical review E, vol. 64, no. 2, p. 026118, 2001.

[21] M. Girvan and M. E. Newman, “Community structure in social and biological networks,” Proceedings of the national academy of sciences, vol. 99, no. 12, pp. 7821–7826, 2002.

[22] V. Batagelj and A. Mrvar, “Pajekanalysis and visualization of large networks,” in International Symposium on Graph Drawing. Springer, 2001, pp. 477–478.

[23] J. A. Grochow and M. Kellis, “Network motif discovery using subgraph enumeration and symmetry-breaking,” in Annual International Conference on Research in Computational Molecular Biology. Springer, 2007, pp. 92–106.

[24] S. Schbath, V. Lacroix, and M.-F. Sagot, “Assessing the exceptionality of coloured motifs in networks,” EURASIP Journal on Bioinformatics and Systems Biology, vol. 2009, no. 1, p. 616234, 2008.

[25] S. Panni and S. E. Rombo, “Searching for repetitions in biological networks: methods, resources and tools,” Briefings in bioinformatics, vol. 16, no. 1, pp. 118–136, 2013.

[26] S. Itzkovitz, R. Levitt, N. Kashtan, R. Milo, M. Itzkovitz, and U. Alon, “Coarse-graining and self-dissimilarity of complex networks,” Physical Review E, vol. 71, no. 1, p. 016127, 2005.

[27] S. Khakabimamaghani, I. Sharafuddin, N. Dichter, I. Koch, and A. Masoudi-Nejad, “Quatexelero: an accelerated exact network motif detection algorithm,” PloS one, vol. 8, no. 7, p. e68073, 2013.

[28] J. Leskovec and R. Sosič, “Snap: A general-purpose network analysis and graph-mining library,” ACM Transactions on Intelligent Systems and Technology (TIST), vol. 8, no. 1, p. 1, 2016.

[29] Y. Chi, Y. Yang, and R. R. Muntz, “Canonical forms for labelled trees and their applications in frequent subtree mining,” Knowledge and Information Systems, vol. 8, no. 2, pp. 203–234, 2005.

[30] D. Bu, Y. Zhao, L. Cai, H. Xue, X. Zhu, H. Lu, J. Zhang, S. Sun, L. Ling, N. Zhang et al., “Topological structure analysis of the protein protein interaction network in budding yeast,” Nucleic acids research, vol. 31, no. 9, pp. 2443–2450, 2003.

[31] T. Rolland, M. Taşan, B. Charloteaux, S. J. Pevzner, Q. Zhong, N. Sahni, S. Yi, I. Lemmens, C. Fontanillo, R. Mosca et al., “A proteome-scale map of the human interactome network,” Cell, vol. 159, no. 5, pp. 1212–1226, 2014.

[32] D. Szklarczyk, J. H. Morris, H. Cook, M. Kuhn, S. Wyder, M. Simonovic, A. Santos, N. T. Doncheva, A. Roth, P. Bork et al., “The string database in 2017: quality-controlled protein-protein association networks, made broadly accessible,” Nucleic acids research, p. gkw937, 2016.

[33] Z. Altun and D. Hall, “Wormatlas,” URL http://www.wormatlas.org, vol. 1384, 2002.

